# Creative destruction: a basic computational model of cortical layer formation

**DOI:** 10.1101/2020.01.29.921999

**Authors:** Roman Bauer, Gavin J Clowry, Marcus Kaiser

## Abstract

One of the most characteristic properties of many vertebrate neural systems is the layered organization of different cell types. This cytoarchitecture exists in the cortex, the retina, the hippocampus and many other parts of the central nervous system. The developmental mechanisms of neural layer formation have been subject to substantial experimental efforts. Here, we provide a general computational model for cortical layer formation in 3D physical space. We show that this multi-scale, agent-based model comprising two distinct stages of apoptosis, can account for the wide range of neuronal numbers encountered in different cortical areas and species. Our results demonstrate the phenotypic richness of a basic state diagram structure, and suggest a novel function for apoptosis. Moreover, slightly changed gene regulatory dynamics recapitulate characteristic properties observed in neurodevelopmental diseases. Overall, we propose a novel computational model using gene-type rules, exhibiting many characteristics of normal and pathological cortical development.

## Introduction

### Background

Biological tissue often exhibits complex but general structures. One well-known example is the layered distribution of different cell types seen in the mammalian neocortex, which has been a central research topic in neuroscience. A number of experimental studies have led to insights into the well-orchestrated processes that form the characteristic layers seen in the adult (Molyneaux et al. 2007; Gaspard 2011). While the pioneer of modern neuroscience Ramon y Cajal and others postulated seven layers in most neocortical regions (Kuhlenbeck 1967), French neurologist Jules Baillarger was the first to postulate six cortical layers (Baillarger 1840) which was later adopted by Korbinian Brodmann in (Brodmann 1909).

Experimental work has demonstrated that the synaptic connectivity and electrical activity dynamics of cortex are strongly shaped by its layer architecture. Along those lines, the main prevailing model of information processing within a cortical column tightly links its functionality to the laminar architecture (Douglas et al. 1989; Douglas and Martin 2004). Also on a global scale do interareal connections obey generic rules dependent upon laminar identity (Markov et al. 2014). Even though it is possible that the layer architecture is not crucial for higher-order cognition, its omnipresence across different species and neural systems suggests it to be a critical factor for neural computation (Striedter 2005; Adesnik and Naka 2018). Along those lines, a number of experimental studies established cortical layer-specific functionalities (Bortone et al. 2014; Lur et al. 2016; Quiquempoix et al. 2018).

To this day, the question of how genetic dynamics give rise to the cortical neural layer architecture remains a key matter of investigation in neuroscience. Already Wilhelm His and Ramon y Cajal studied how proliferating cells in the ventricular zone (VZ) generate neurons migrating radially towards the pia and stop before the marginal zone (MZ) (y Cajal 1891; His 1904). A better understanding of how layers form during cortical development is relevant from a biological as well as medical point of view. In particular, many brain pathologies, such as in autism (Crawley 2012), schizophrenia (Nawa et al. 2000), Fetal Alcohol Spectrum Disorder (FASD) (Riley et al. 2011) or epilepsy (Bozzi et al. 2012) have been associated with developmental origins of pathological layer formation.

In recent years, *in vitro* models have become a viable approach to conduct research on neural development (Eiraku and Sasai 2012; Lancaster et al. 2013; Lancaster and Knoblich 2014). However, despite rapid progress along those lines, a number of serious challenges in terms of recapitulating *in vivo* conditions and functional behaviour remain. A complementary, computational approach can benefit this research direction in multiple ways; it enables to combine various data into a coherent framework, and allows to formulate and assess novel hypotheses in a quantitative manner. In particular, mechanistic computational models constitute a promising approach, because they facilitate the incorporation of various genetic, molecular and imaging data, and so constitute a strong link between experiment and theory.

Only few computational/mathematical studies provide a quantitative framework for cortical layer formation. Cahalane et al. show that a mathematical model of the dynamics of neural fate determination can yield anatomical properties of superficial and deep layers of cortex across different species (Cahalane et al. 2014). However, since this framework is formulated on an abstract, analytical level, it does not take into account mechanical interactions between cells, or the setting within the spatial environment. Hence, it is problematic to test more direct, mechanistic hypotheses in such an approach, e.g. on the involvement of certain pathways or extracellular cues. A different computational approach is taken by Caffrey et al., who employ an agent-based model to model cortical layer formation (Caffrey et al. 2014). This model does not comprise the developmentally crucial aspect of proliferation, and relies on highly simplified intercellular interactions. Along those lines, other computational studies that model axonal and dendritic trees do not focus on the origins of a layered architecture (Koene et al. 2009; Cuntz et al. 2010). In contrast, Zubler et al. present an agent-based computational model for cortical lamination with mechanical interactions (Zubler et al. 2013). Based on this approach and software (Zubler 2009), cortical development has been studied from a conceptual perspective of self-construction and put in correspondence with genetic data (Zubler et al. 2013). Importantly, this approach relies solely on modelling local information exchange, hence it is maintained the biologically realistic condition that cells can only experience their local 3D environment.

Here, we build upon the approach of Zubler et al. (2013). Going beyond a qualitative assessment, we here provide a computational model that reproduces quantitative measurements of the cortical layer architecture. In particular, we study how the wide range of neocortical layer architectures observed in health and disease, across species and cortical regions, arises. To this end, we devise a canonical gene regulatory network (GRN) model, which individually plays out from an homogeneous pool of progenitor cells. In other words, we study how a core GRN, by differential alterations to its physiological interactions (Macneil and Walhout 2011), can generate a wide range of cortical layer architectures. In order to account for the evolutionary conservation of this basic structure, we reproduce experimental measurements from the human, macaque, rat and mouse cortex. In particular, we investigate how cortical layers comprising different numbers of neurons can arise and evolve in a biologically plausible way. Importantly, this GRN model comprises the key ingredients of cortical development, i.e. it incorporates the interplay between neuronal proliferation, differentiation and apoptosis. Moreover, it respects detailed characteristics of cortical layer formation, such as an initial exponential proliferation phase that is followed by a sequential differentiation phase (Lui et al. 2011).

The overall computational model is tested in a so-called agent-based approach: initially, a homogeneous pool of neuronal precursors is represented, where each cell’s behaviour is guided by its individual gene regulatory network model, as predicted by the protomap hypothesis (Rakic et al. 2009). This model constitutes the genetic rules that specify the cellular dynamics of the individual cells (hence each cell acts as an autonomous “agent”). Based on this initial configuration, simulations are conducted to demonstrate how, from the proliferating cell population, the genetically specified dynamics play out and yield laminar neuron numbers in accordance with experimental observations.

## Materials and Methods

### Materials

#### Layer-specific neuron numbers

Experimentally obtained measurements of layer-and species-specific neuron numbers were obtained from (O’Kusky and Colonnier 1982; Schuz and Gunther 1989; DeFelipe et al. 2002). Layer thicknesses from human cortex were obtained from (Economo 2009).

#### Images

The histology sections shown in Fig. 7 were obtained from (Casanova 2007) and (Golden and Harding 2010).

### Methods

#### Simulation environment

The simulations were conducted using the open-source framework Cx3Dp, the parallelized version of Cx3D (Zubler 2009) (available at http://www.ini.uzh.ch/projects/cx3d/). We implemented an agent-based model using a parallelized version of Cx3D.

Our computational model specifies a state machine that defines the network of cell state transitions that can be understood as an abstract GRN. It is instantiated in a few precursor cells at the beginning of the simulation. Each of these cells follows internal, gene-type rules, while physically interacting with the local, extracellular environment. Mechanical forces also act between neighboring cells that are in physical contact. These cell-cell interaction forces are computed based on the diameter of spherical somata, their relative distance, and their adhesive properties (for more information on the computation of the mechanical forces, please refer to (Zubler 2009)). Moreover, cells can secrete chemicals that diffuse extracellularly in 3D space in a gradient fashion, and that can be sensed by other cells.

#### Experimental data

Layer-specific neuron densities in different cortial areas in different species were obtained from (O’Kusky and Colonnier 1982; Schuz and Palm 1989; DeFelipe et al. 2002). In order to compare with the computer simulation results, the number of neurons per layer under a cortical surface area of 300μm×300μm was computed. Moreover, since these numbers reflect the neuronal statistics in the adult cortex, the number of neurons dying due to activity-dependent apoptosis (i.e. apoptosis influenced by synaptic connectivity and electrical activity, which was not included in our study), estimated to amount to 25% of all neurons (Heck et al. 2008), was added to the adult-stage neuron numbers. Hence, the numbers listed in Supplementary Table S1 are multiplied by 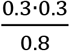 to compare with the computer simulation results.

Layer thicknesses used to inform the phenomenological model were obtained from (Economo 2009). Time stamps in Fig. 3 are based on (Saito et al. 2011; Workman et al. 2013).

#### Simulation setting

Initially, 175 precursor cells are arranged as a 2D layer comprising an area of 300μm×300μm, where the cell body locations are given by (normally distributed) random displacements from regular gridpoints with a jitter of 3μm standard deviation. The simulation takes place within a 3D environment with columnar boundaries of size 300μm×300μm×4500μm, the latter being the radial dimension.

Standard Cx3Dp cell body parameters are chosen for these. Each precursor cell is initialized with an intracellular substance quantity of 100 (unitless), which decays by 1 % at each time step. The end of the initial, symmetric proliferation phase is invoked when this substance quantity falls below a prespecified threshold Tsym. 50,000 times steps were simulated to generate the final laminar arrangement. The model produced systems containing approximately 7,000, 11,000, 16,000 and 24,000 cell bodies in human, rat, mouse and macaque cortex simulations, respectively.

#### Model parameters

The maximum cell body diameter during proliferation and differentiation was set to 10 μm. If this value is reached during cell body growth, the cell body either divides and subsequently increases its diameter during growth (the diameter growth per time step was 0.6 μm), or terminates its differentiation pathway. However, after a cell has terminated migration and has taken up its final location within the layer architecture, it grows its diameter to 15 μm, expressing its mature phenotype.

Certain physical parameters change dynamically during the simulation, in order to improve the segregation between the individual layers. In particular, marginal zone cell bodies that have reached their pre-specified external substance concentration and stopped their migratory mode, adopt the Cx3D parameters *interobject-force* coefficient of 1.1, *adherence* of 0.00001 and a *mass* value of 0.00001. The slightly higher than normal interobject-force coefficient (standard value 1) renders the marginal zone cells more responsive to mechanical forces from migrating cells. Also, the small adherence and mass values enable them to be easily pushed, for if their mass were large they would remain at their original position.

The state machine model of the GRN has 12 parameters. These comprise six probabilities that specify the probability to commit to layer-specific neuron types (C_M_, C_2_, C_3_, C_4_, C_5_, C_6_). Moreover, the five parameters P_M_, P_2_, P_3_, P_4_, P_5_ and P_6_ determine the probability to commit to apoptosis of type A1. For simplicity, C2+P2=1 was used, so only 11 of the 12 parameters were varied to match the experimental data. The (fixed) parameter Tsym determines the number of symmetric cell divisions in the early proliferation phase, and is set identical for all the layer architectures here for the sake of simplicity.

Marginal zone cells that have terminated migration each create a physical bond with a neighboring cell, with the Cx3D physical bond parameters *breaking-point* set to 10,000%, the *dumping constant* set to 0, and the *spring constant* set to 1. This physical bonding enables marginal zone cells to remain connected with one another. In contrast, cell bodies of the other layers (2-6) assume a mass of 0.1 during migration, and increase their mass to 10 after termination of migration. Moreover, migratory cells’ *interobject-force* coefficients assume a value of 0.0001. Overall, these physical parameter settings facilitate the right developmental operation: heavy cells push the lighter marginal zone cells before taking up their final position. Preliminary simulations indicate that limited variations of these parameters do not significantly alter the resulting cell distributions.

The stopping of cell migration is induced when certain conditions are fulfilled. In the case of layer 2, 3, 4 and 5 cells, two conditions need to be fulfilled: the number of marginal zone cells in the local neighborhood, i.e. in the physical space where physical interactions can take place, must exceed one. Moreover, the number of neighboring cells of the other, earlier developing layers (e.g. in the case of layer 4 cells, these would be layers 5 and 6) must be zero. In the case of layer 6 cells, at least one marginal zone cell must be detected. Finally, in the case of marginal zone cells, a pre-specified extracellular cue concentration needs to be reached. To this end, an extracellular substance gradient that increases in concentration towards the pia, is instantiated when stimulations start. These conditions enabled cells to recognise their appropriate locations, and establish the associated layer thicknesses while pushing the marginal zone towards the pia.

Once cells have stopped, they initiate their apoptosis modules that trigger cell death if a given condition is fulfilled. Neurons belonging to a specific layer sense the number of local, neighboring neurons (distance < 20 μm) belonging to a different layer. If at least three neurons belong to neighboring layers, or one belongs to a more distant layer (e.g. layer 6 for neurons in layer 3), this condition is fulfilled, and (A2-type) apoptosis is triggered.

Since the number of layer 1 cells is very low in all species, an additional probabilistic apoptotic rate was included only for layer 1 cells, which does not depend on extracellular conditions. This apoptotic probability was set to p=0.25, 0.9, 0.2 and 0.8 for human, rat, mouse and macaque cortex, respectively, which yielded agreement with the experimentally measured neuron numbers in layer 1.

#### Analysis of layer properties

After simulations terminated, the cell positions and information on their cell identity (i.e. which layers they belong to) were exported as a matrix into a Matlab-readable file. Since (occasionally) cells migrated through the marginal zone and ended up at the topmost radial position, these were discarded in the analysis by thresholding.

Simulated layer arrangements were measured using the Matlab *ksdensity* function. This function computes the normalized distribution of individual layers. This distribution was then multiplied by the number of total cells in a layer, which yielded a smooth distribution of cells across the radial, spatial dimension.

The histograms shown in Fig. 4 were also computed using Matlab. In order to better account for the variation of layer-specific cell numbers, the number of bins for any given layer was chosen as 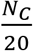, where NC is the total number of cells of this layer. Hence, cell distributions within larger layers are displayed using more bins.

The t-score and t-statistics for the various layer-specific neuron numbers that are compared in Fig. 5 were computed using Matlab. A two-sided t-test with significance level α = 0.05 was conducted.

## Results

### The model

We studied two phases during cortical development, i.e. the progenitor amplification phase, when the neural progenitor cell pool increases exponentially due to recursive cell division. In the second phase, differentiation of neurons and migration is modelled. Fig. 1A visualises these two phases, which occur in succession.

**Figure 1:**
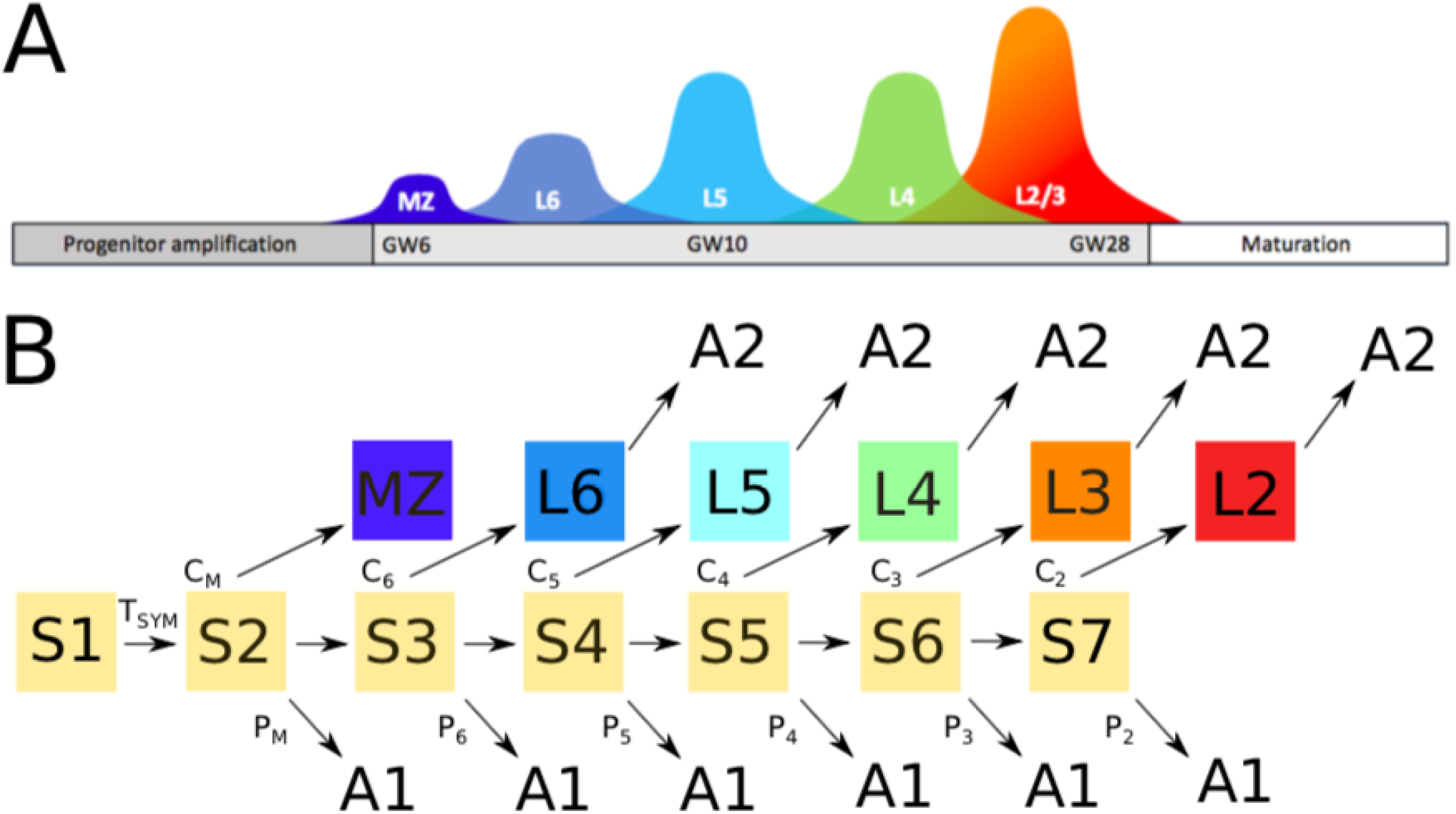
(A) Schematic of sequence of cortical layer formation. During cortical development, progenitor cells proliferate and differentiate into different neuron types in a consistent sequence. Marginal zone (MZ) cells (dark blue) are the earliest cells to be produced. Afterwards, differentiation into layer 6 (blue), layer 5 (cyan), layer 4 (green), layer 2/3 (purple) cells occurs. (B) Structure of state diagram of cortical lamination model. Arrows indicate (probabilistic) paths of the network constituting the state diagram. The initial, purely proliferative state S1 leads to state S2 when an internal timer triggers this state transition. From this second proliferative state S2, neuronal migration of MZ cells is triggered with probability C_M_ (the marginal zone is the predecessor of layer 1). Alternatively, apoptosis (A1-type) is invoked, or the cell divides again to proceed to the next neural progenitor state S3. S3 cells are the progenitors of layer 6, layer 5, layer 4 or layer 2/3 cells. During this sequential differentiation process, apoptosis occurs with probabilities P_6_ to P_2_. At the end of the lamination process, a final apoptotic step (A2) occurs that depends on the local neighborhood of cells. Notably, the final neural precursor state S7 does not yield any cell division, and so the condition C_2_ + P_2_ = 1 is satisfied. Yellow colour indicates stem cell states, i.e. these cells have the potential to divide. Dark blue, light blue, cyan, green, orange and red indicate final cell types.

#### The progenitor amplification phase

The simulation starts with the progenitor cells that are arranged randomly on a 2D sheet, which represents the proliferative zone. Each of these cells follow their own gene regulatory dynamics, specifying recursive cell growth and division (Fig. 1B). The initial state of the GRN is S1. In this first stage of proliferation, cells divide and grow recursively, producing an exponentially increasing number of neural progenitor cells (S2) (Lui et al. 2011). Squares coloured yellow indicate cell states with proliferation capacity.

In our model, the proliferating cells possess an internal clock that is realized by means of an intracellular substance. This substance decays over time, and once the progenitor cells sense that it is below a prespecified threshold T0, they terminate this first proliferative stage of symmetric division. These progenitor cells then transition to stage S2, which is the first GRN state that leads to neuronal differentiation. Hence, this intracellular substance acts as a trigger for the neuron specification phase.

#### The differentiation phase

After the exponential proliferation phase (instantiated within the cell state S1), the progenitor cells still retain the potential to proliferate, but they additionally can probabilistically commit to various other cell types (Gao et al. 2014). Once this differentiation process has occurred, neurons migrate radially to take up their final location within the cortex.

Initially, the neuronal progenitor cells (probabilistically) either differentiate into marginal zone (MZ) cells, progress further along the differentiation path by dividing, or undergo apoptosis. States S2 to S7 constitute proliferative stem cell states, where (for simplicity) only one round of proliferation can occur.

The MZ develops by MZ cells migrating along the gradient of a predefined extracellular cue that increases in concentration along the radial direction. The MZ cells continuously sense this chemical cue, and once a given threshold is reached, they stop migrating. The layer structure is not influenced by the magnitude of this threshold, which solely determines the distance between the proliferative zone and the developing cortex. It mimics the influence of the pial membrane, which in reality stop migration of the MZ cells.

For simplification, we model that layer 6 migrate separately from MZ cells, although in reality layer 6 splits the so-called preplate into MZ and subplate. In our model, layer 6 cells also commit to migration, after a brief waiting period (that is specified again based on a decaying intracellular substance acting as a cellular clock). Because of the waiting period, L6 cells migration occurs after MZ cells have reached their target, and L6 cells stop migrating as soon as they sense MZ cells.

The rest of the differentiation process is a repetition of this elemental process of differentiation, proliferation and apoptosis: those precursor cells that did not commit to the MZ or L6 fate will continue to divide. After dividing, they again probabilistically commit to the neuronal fate (i.e. layer 5), continue with proliferation, or undergo apoptosis, and so on. This developmental process can be very efficiently encoded in the GRN, because it is composed of repetitions of the same elemental genetic rule.

The parameters of the GRN are given by: Tsym, which determines the number of cell divisions in the progenitor amplification phase; C_M_, C_6_, C_5_, C_4_, C_3_ and C_2_ which determine the probabilities for progenitor cells to commit to the associated, differentiated neuron types; P_M_, P_6_, P_5_, P_4_, P_3_ and P_2_, which are the probabilities for progenitor cells to commit to apoptosis.

### Consistency of gene-regulatory network structure

One remarkable observation across different cortical areas in humans as well as in other species is the significant variation of cell numbers in individual layers (Fig. 2) (Herculano-Houzel et al. 2008; Collins 2011). It follows that an important condition for a mechanistic generic computational model of layer formation is the possibility to account for various compositions of layer-dependent cell numbers.

**Figure 2:**
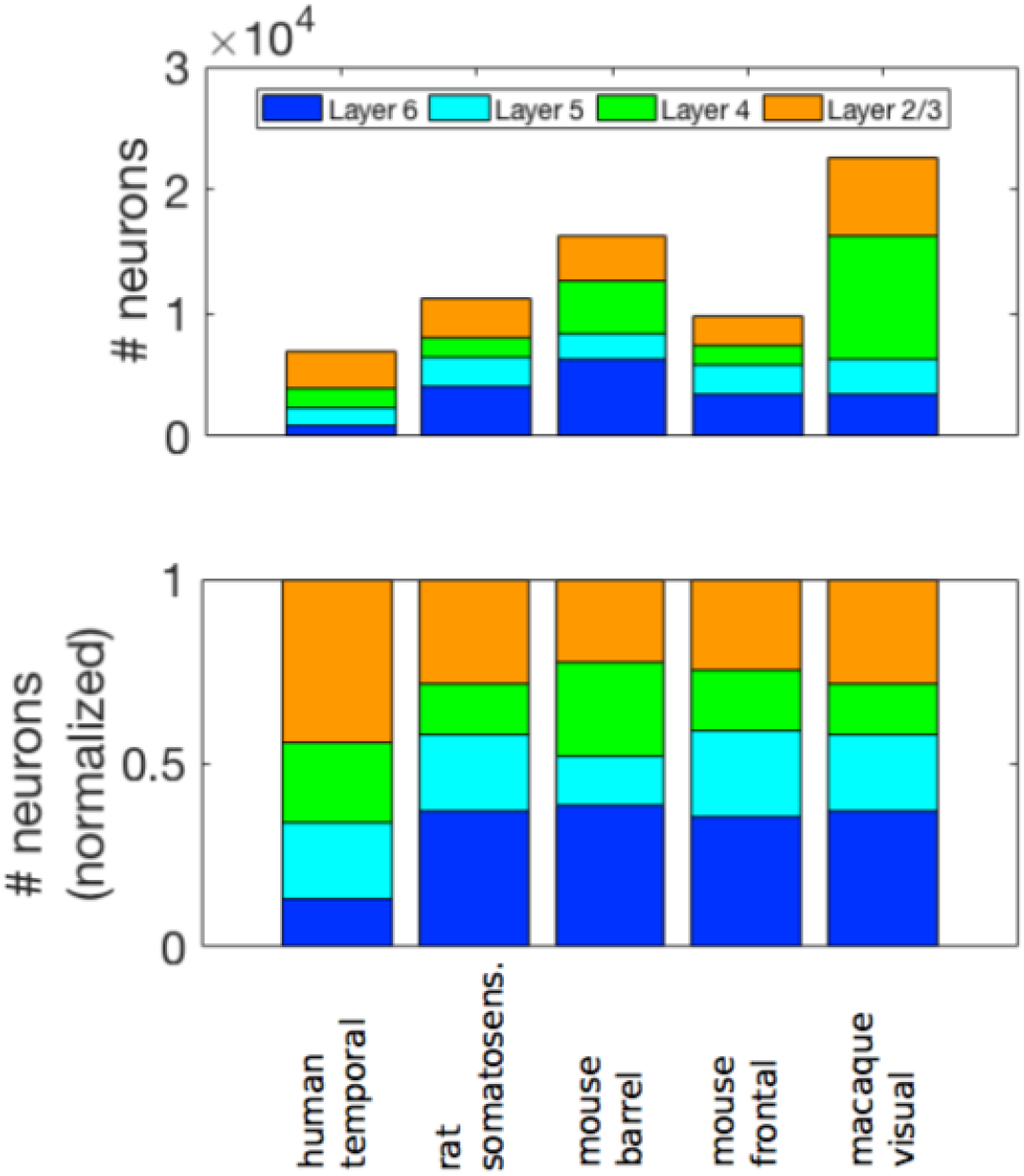
Number of neurons under 1 mm2 of cortical surface area, in the cortical layers of different areas and species. The shown examples are: human temporal cortex, rat somatosensory cortex, mouse barrel cortex, mouse frontal cortex and macaque visual cortex. There is substantial variation in the dimensions of individual layers (A), and the relative proportions (B) within the same cortical area and species. This variation in cortical layer thicknesses stems from differences in neuronal size, as well as cell numbers.

We first demonstrate that our model can yield the neuronal numbers across layers within different cortical areas. Fig. 3 shows the sequential development of the layers in the human temporal cortex in the correct order, i.e. the characteristic inside-out dynamics of layer formation is recapitulated (see also Suppl. Video 1). Moreover, the development of the layers in rat, mouse and macaque cortices was simulated again by introducing changes to the model parameters. The resulting distributions of neurons across different layers are shown in Fig. 4. The normalised probability distributions of these neuronal locations are shown in Suppl. Fig. S1.

**Figure 3:**
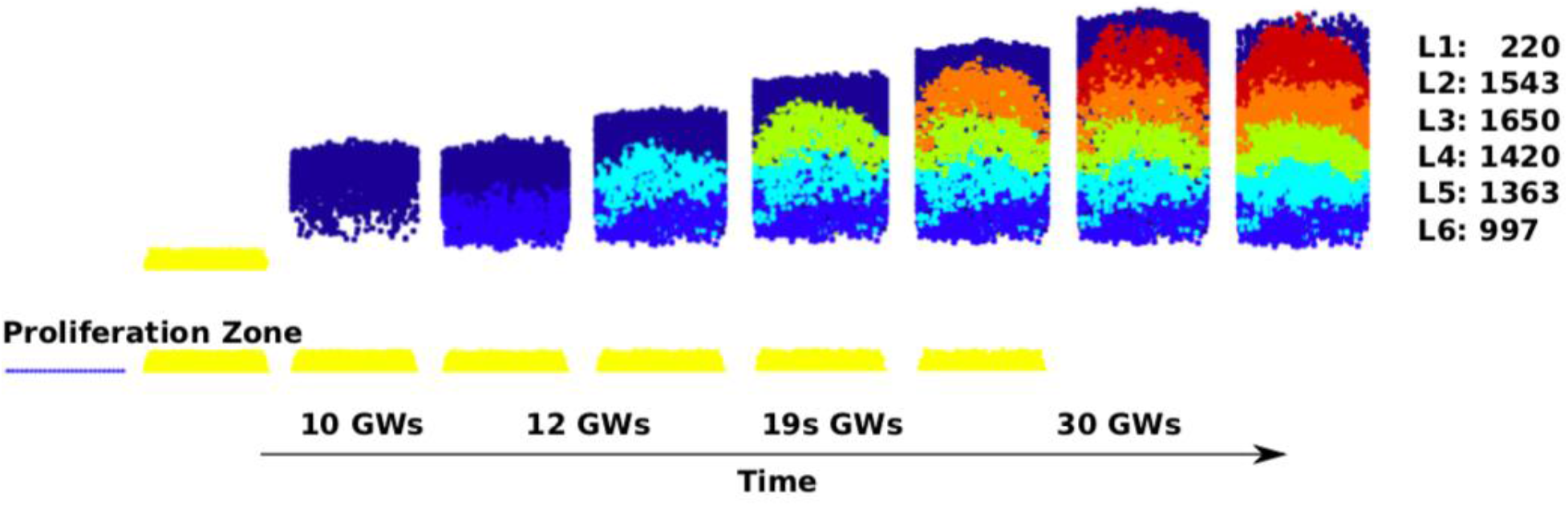
Sequence of lamination steps. In this simulation of layer formation in the peristriate area of the human temporal cortex, initially only a small set of progenitor cells exist in the ventricular zone (left, dark blue). This proliferative zone (yellow) generates neuronal precursors in an exponential manner. These precursors, via symmetric and asymmetric division, give rise to different cortical layers. The first cells that migrate are marginal zone cells (dark blue, top). Subsequently, layer 6 cells (blue) physically push the marginal zone, constituting a mechanistic process that is repeated for the layers 5 (cyan), 4 (green), 3 (orange) and 2 (red). Hence, the layer architecture is produced in the characteristic outside-in manner of cortical layer formation. In humans, the overall formation of cerebral cortex happens approximately between gestational weeks 8 and 30.

**Figure 4:**
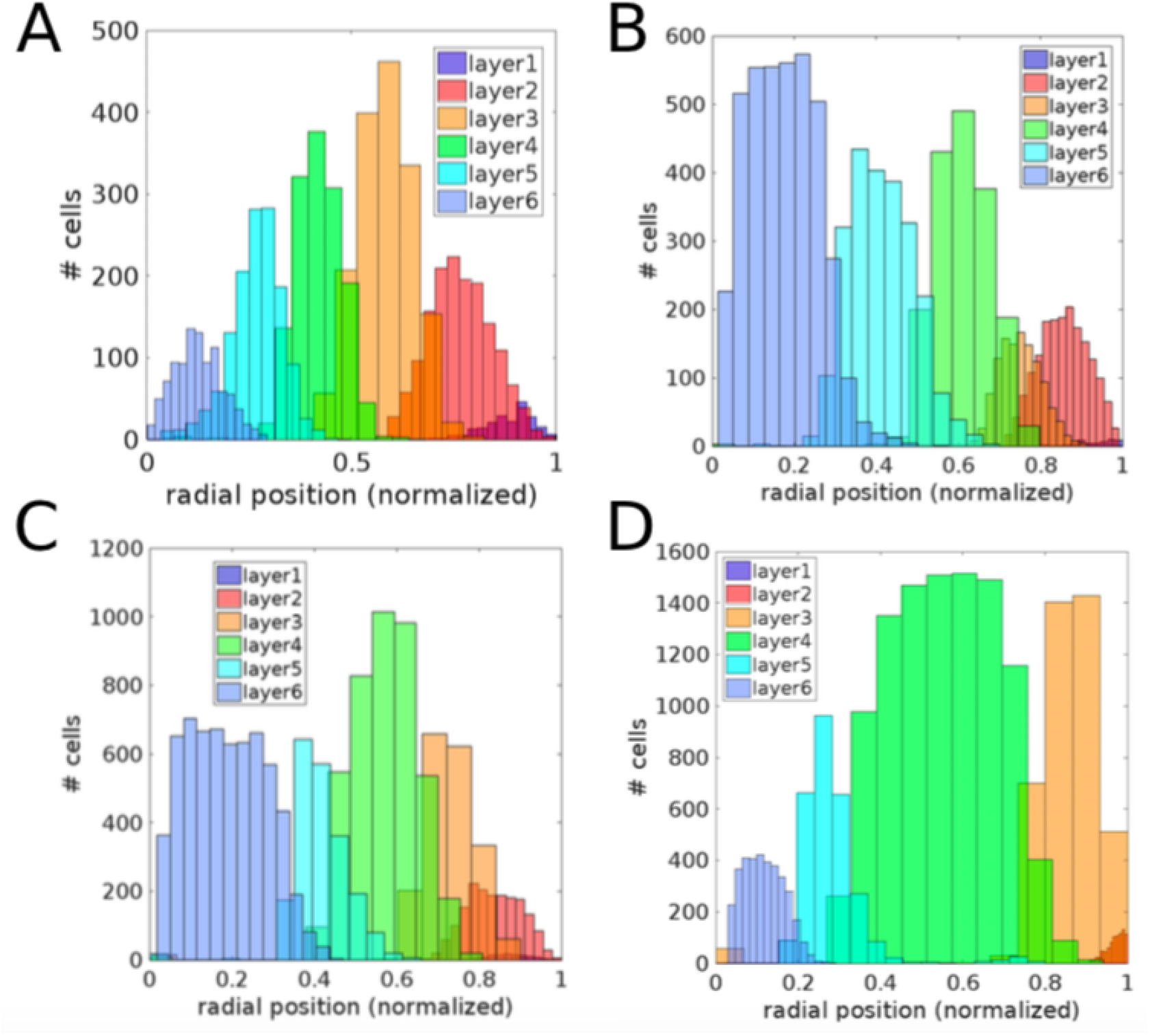
Spatial distributions of simulated, layer-specific neuronal locations along the radial axis. The shown layer architectures are from (A) human temporal cortex, (B) rat somatosensory cortex, (C) mouse barrel cortex and (D) macaque visual cortex.

In Fig. 5, the experimental data obtained from multiple experimental studies are shown together with the corresponding simulated layer dimensions, highlighting the explanatory power of the simple GRN structure shown in Fig. 1. Overall, these results demonstrate that inter-species variation of layer-specific cell numbers can be accounted for only by modifications to the parameters of the GRN, rather than structural changes.

**Figure 5:**
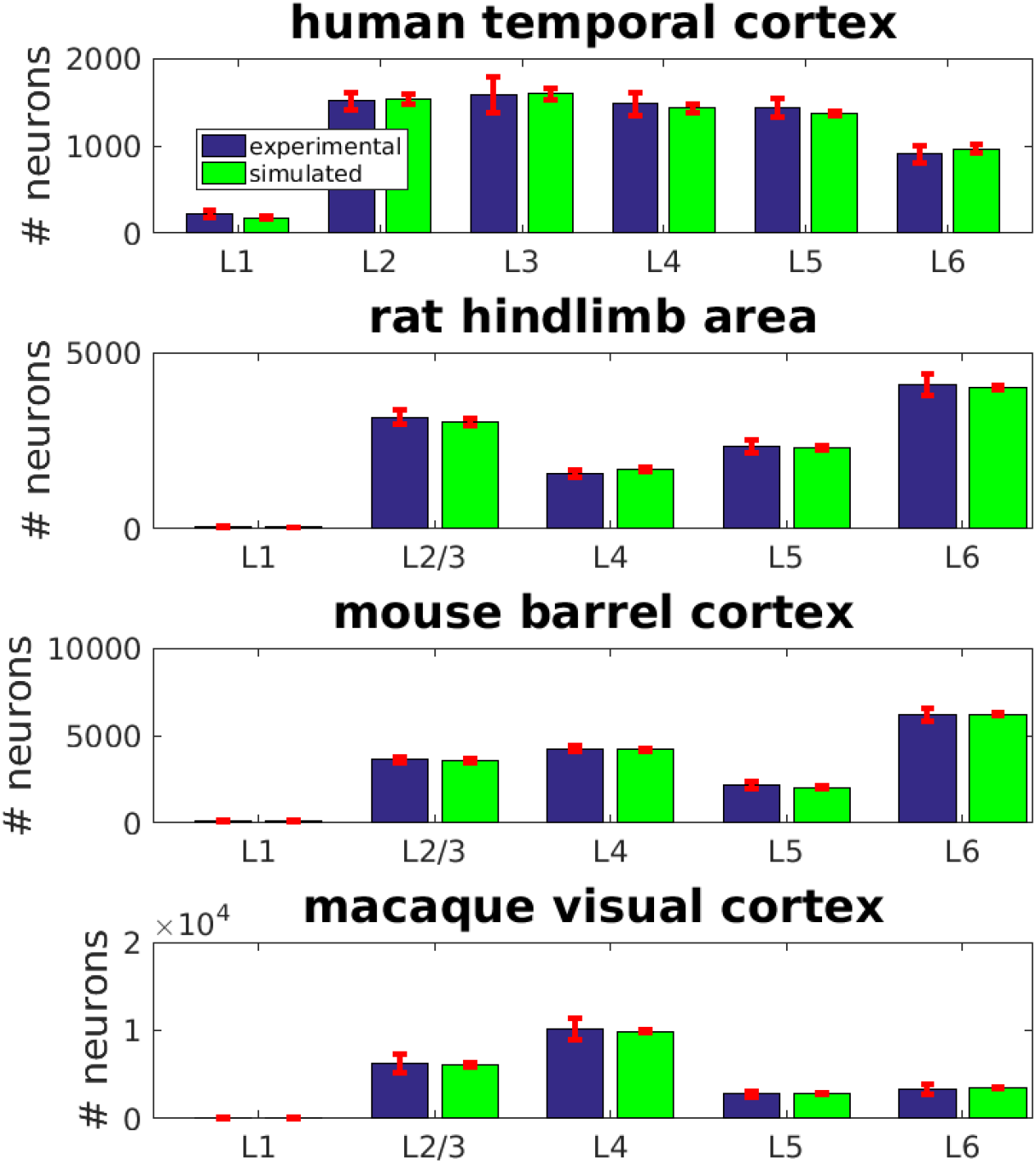
Layer-specific neuron numbers in human, rat, mouse and macaque cortical tissue, under one square millimetre of cortical surface. Numbers based on experimentally measurements and based on simulations are shown in blue and green, respectively. The simulated layer architectures are in accordance with the experimentally obtained measurements, demonstrating that the model can consistently account for a wide range of neuron numbers. Error bars (red) indicate standard deviation, as obtained from experimental data and from 10 simulations, respectively. The null hypothesis that the means of the simulated layer-specific neuron numbers stem from the experimentally derived numbers could not be rejected based on a two-sided t-test with significance level α = 0.05, for all the shown data.

Fig. 6 shows the cell number dynamics in a simulation of human cortical layer formation. First, the marginal zone that will subsequently become layer 1 develops, and then layers 6, 5, 4, 3 and 2 form. The initial production from the stem cell pool is followed by reductions in cell numbers through the apoptotic stage A2 (Fig. 1). We find that most of the apoptosis occurs in mislocated MZ cells, predominantly in the superficial cortical layers.

**Figure 6:**
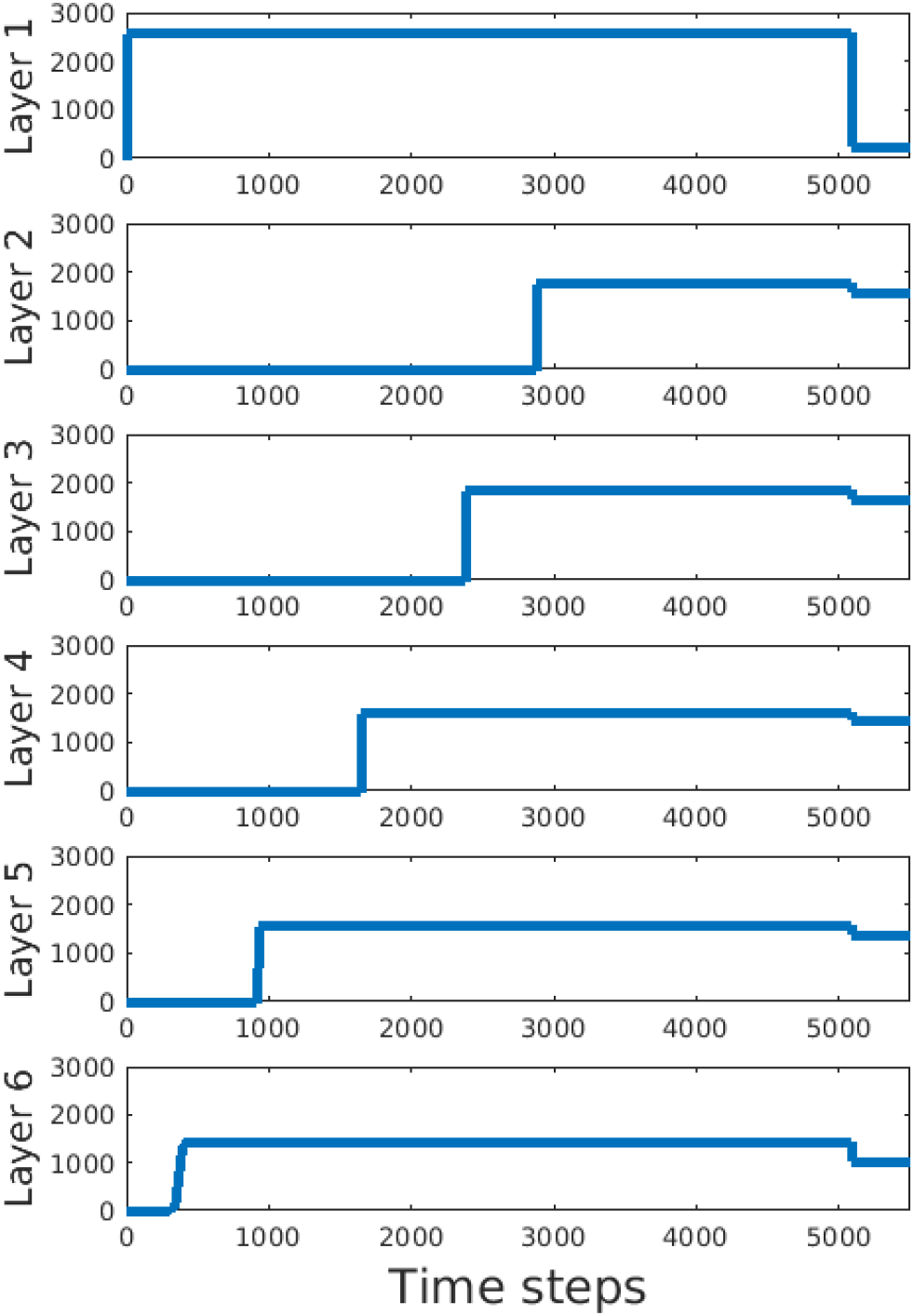
Neuron numbers in individual layers during layer formation. The GRN defines a sequential differentiation process, whereby the individual layers originate in a predefined temporal order from the proliferative zone. This developmental sequence expresses itself by a rise in neuron numbers, whereby first the marginal zone (later constituting layer 1) develops. Subsequently, the deep layers form before the superficial layers. Finally, neurons that recognize their own mislocation by identifying their immediate neighbour neuron types, enter the apoptotic state A2, which yields a decline in neuron numbers. In our model, this apoptotic state is triggered by layer 2 cells that occupy their terminal location after migration and signal this event to the other cells via an extracellularly diffusing substance. Note the large proportion of apoptosis for layer 1 cells, which stems from the fact that these cells are pushed by the later-developing neurons, and so exhibit a higher risk for mislocation.

The modifications of the GRN are solely changes in probabilities for transitions between certain states, rather than changes to the GRN states or connections themselves. This general applicability of the GRN structure indicates that minor evolutionary changes can yield a wide range of phenotypic differences in brain structure. In our model, these modifications can include changes to apoptotic rates, in addition to changes to cellular fate commitment. However, these model parameters are under-constrained by current experimental data, hence there are different possibilities of GRN parameter sets yielding equivalent layer-specific cell numbers. Suppl. Fig. S2 shows how changes in exemplary parameters of the GRN affect the resulting layer-specific neuron numbers: depending on where in the GRN the given model parameter takes effect, the subsequently developing layers are affected.

### Differential apoptotic processes

In our model, changes to apoptosis at early stages affect the final cell numbers. Apoptosis later during development has less impact on the total tissue thickness, because only later developing layers will be affected. Nevertheless, all the probabilities for cells in the proliferative stages to exit and commit to apoptosis (GRN states labelled A1 in Fig. 1) exert control over cell numbers. Overall, changes to these probabilities can yield extensive differences in the final cell numbers and layer dimensions, and so the A1-type apoptotic stage is paramount in generating the right number of cells.

In a second process, apoptosis is a crucial determinant of the proper layer architecture and neuronal density (GRN states labelled A2 in Fig. 1). Our model of this second type of apoptosis is inspired by (Hauri 2013): cells can sense their local environment, and based on the neighboring cell types commit to apoptosis. Whenever a cell senses too many neurons of a type that is different to its own, it commits to apoptosis. Hence, in this second phase of apoptosis, a different pathway is activated, because it involves the binding of extracellular ligands (Guerin et al. 2002).

In our model, this A2-type apoptosis is initiated after the end of migration, when the first layer 2 cells have reached their final radial position, and then trigger the apoptotic process. To this end, these apoptosis-initiating layer 2 cells secrete a diffusible substance, which is then detected by all cells that have reached their final destination. Once a sufficient amount of this factor is detected in their neighborhood, these cells sense the number of cells different from their own type. If this number exceeds a prespecified threshold, the cell commits to apoptosis. This type of apoptosis that happens after migration in our model is crucial for refining the layered architecture of the final cortical structure.

These simulation results point towards apoptosis as a highly heterogeneous mechanism. In the first apoptotic stage, the fate is determined probabilistically, according to a pre-specified chance that is not influenced by external factors. In the second apoptotic stage, extracellular cues conveying information on the identities of neighboring cells are taken into account. In our simulations we observe that this apoptotic behaviour is particularly important for layer 1, which during development is successively pushed by the deeper layers; it occurs more often for this layer that some cells are not pushed effectively, hence remain stuck in a deeper position than the target layer.

### Two-factor based determination of layer-specific neuron numbers

In our model, the number of neurons per layer is determined by overall 11 GRN parameters. These parameters specify the probabilities to commit to either a certain layer neuron type, continue proliferation and then potentially apoptosis. These parameters are C_M_, C_6_, C_5_, C_4_, C_3_, C_2_, P_M_, P_6_, P_5_, P_4_ and P_3_ (P_2_ is given by 1-C_2_). Based on these parameters, we derived the number of neurons per layer (the GRN is inherently stochastic and so we estimate the expected number of neurons).

We indicate the number of undifferentiated precursor cells at state S2, after the symmetric division phase has ended, by NP. The number of cells in layer 1 is then proportional to NP and the probability to commit to the marginal zone cell fate (*C*_*M*_):

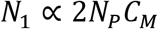

The factor 2 stems from the fact that there is a cell division at state S2. Moreover, the number of layer 6 cells (*N*_6_) is determined by two parameters, namely *P*_*M*_ and *C*_6_. The former has a negative impact on *N*_6_, while the latter increases it. Analogously, *N*_5_ is antagonistically determined by two further parameters, i.e. *P*_6_ and *C*_5_, and so on.

Hence, in our model the layer-specific numbers can change without affecting previous or subsequent layers. In contrast, if there were no apoptosis, any change in a probability to commit to a certain layer would necessarily influence all subsequent layers. For instance, if the probability to commit to layer six would increase, this would mean that the cell numbers in each of layers 5, 4, 3 and 2 would be smaller. Hence, the inclusion of apoptosis enables layers to develop individually, allowing for a high adaptivity of the layer architecture. From an evolutionary perspective, layer architectures can adapt to different requirements in the context of their inputs, outputs and intra-regional circuitry.

### Layer architecture in neurodevelopmental disorders

In addition to layer architectures in health cortex, we also investigated whether our model could capture characteristics of certain neurodevelopmental disorders. Evidently, there is a large variation of the phenotypic expression across pathologies. Nevertheless, specific general properties of malformation and developmental causes are consistent within types of disorders (Barkovich et al. 2005). One example is polymicrogyria, which usually entails a reduced cortical thickness and an increase in folding patterns, while a clearly layered structure is still preserved (Golden and Harding 2010; Judkins et al. 2011). We conducted small changes to the probability parameters of our computational model, leading to an increase in apoptotic activity at early developmental stages (A1). As shown in Fig. 7E, these simulations recapitulate a reduced cortical thickness compared to the control thickness (Suppl. Fig. S3B), while laminar organization is still preserved.

**Figure 7:**
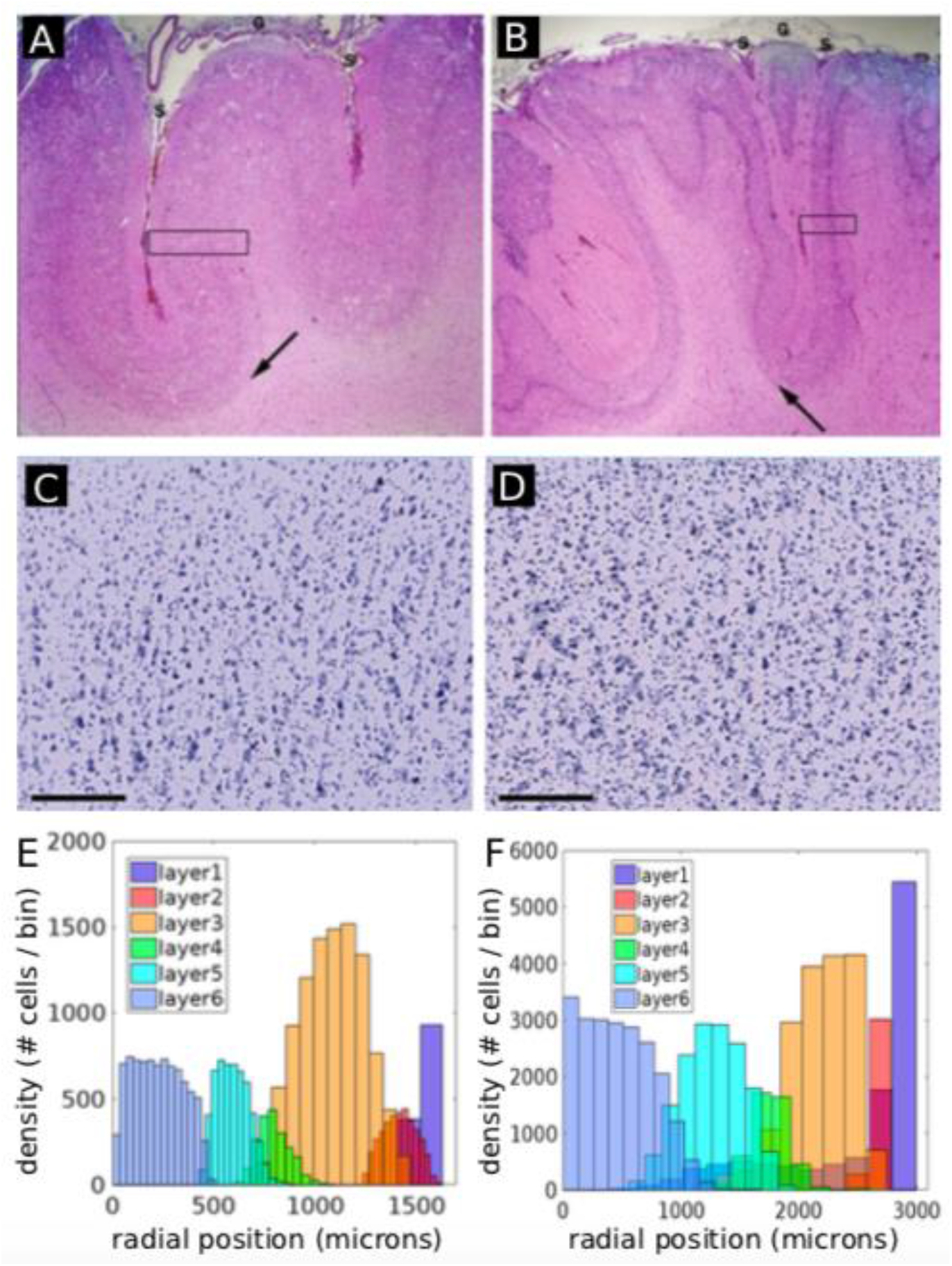
Pathological properties in polymicrogyria and autism can be recapitulated using a phenomenological model of cortical layer formation. (A) Histology section from healthy human cortex and a human cortex with polymicrogyria (B). Patients with polymicrogyria often have a reduced cortical thickness, while more and smaller gyri can be observed. The layer structure is usually preserved. (C, D) Nissl-stain from healthy (C) and autistic (D) cortex (cortical area 9, right hemisphere, layer 3). Experimental observations in the autistic cortex show a higher cell density and more jittered cell arrangement. Scale bars measure 200 μm. (E, F) Intriguingly, we could recapitulate thinner cortices or increased cell density by a single parameter change. (E) Increased (A1-type) apoptosis during early proliferation and differentiation yields simulation results resembling a pathological phenotype, where the cortical thickness is approximately 1.5 mm instead of 3 mm. (F) Dysfunctional apoptosis (A2-type, as indicated in Fig. 1) yields higher cell densities and more mixed positions, reminiscent of observations from the autistic cortex. Especially layer 1 neurons are affected because they are pushed during the entire layer formation process, yielding also cells that are not pushed correctly to the correct position along the radial axis. Without A2-type apoptosis, there is significantly less segregation between neighbouring layers, as indicted by the overlap of the histograms.

In addition to the scenario where apoptosis at early stages was increased, we conducted simulations where the initial phase of proliferation stopped prematurely. This decreased the neural progenitor pool significantly but did not interfere with the migration and apoptosis. As expected, a reduced cortical thickness was observed also in this scenario, while the layer thicknesses were proportionally preserved. Hence, genetic changes reducing the proliferation in the exponential proliferation stage, or increasing A1-type apoptosis, lead to the same result (see Suppl. Fig. S3A).

Also autism spectrum disorders have been associated with pathological apoptosis during brain development (Gabriele et al. 2014). Along those lines, post-mortem histological analysis indicates a higher cell density and less clearly structured tissue organization in terms of cell locations (Casanova 2007). In order to incorporate these observations, we simulated a model where A2-type apoptosis is not functional. As a result, a higher cell density is observed. Moreover, the layers are less clearly separated, and more neurons are mislocated (Fig. 7F). Hence, changes to both types of apoptosis, i.e. either in the first stage or second stage, can yield architectures reminiscent of certain neurodevelopmental disorders.

Finally, we conducted simulations of layer formation in subcortical band heterotopia (also known as double cortex syndrome), where neuronal migration is affected (Guerrini and Dobyns 2014). To this end, differentiated neurons were attributed a malfunctioning migratory behaviour with a 20 % chance. Once differentiated, these pathological cells did not migrate radially, but remained in the proliferative zone. This resulted in the abnormal presence of neurons in the proliferative zone (Suppl. Fig. S4), in agreement with the observations of a deep cellular layer composed of heterotopic neurons (Manent et al. 2009).

## Discussion

The presence of an abundant variety of layered structures is a common property of vertebrate brains. Although a fully comprehensive picture of cortical structure and function is currently lacking, developments in neural imaging techniques have enabled key insights into the intricate relationship between laminar organization and electrical activity (Larkum et al. 2018).

Notably, a mechanistic and in-depth understanding of how cortical layers arise is currently lacking. Computational models are a powerful tool to devise new hypotheses, and to generate experimentally testable predictions. In this work, we provide a model for the development of cortical layer architectures and demonstrate that it is applicable to different cortical areas and species. To this end, our results are in accordance with the observation that the number of cortical neurons varies significantly across the cortical surface (Herculano-Houzel et al. 2008; Collins 2011). Moreover, we show that a number of key characteristics of neurodevelopmental disorders can be accounted for. Our model is based upon apoptosis during development as a highly heterogeneous process, comprising distinct apoptotic stages that depend on intracellular and extracellular conditions, respectively.

Conceptually, our work is different from traditional models of biological, developmental dynamics, in that it follows an agent-based approach. Our model does not rely on any pre-specified neural architecture, is initialized solely from a homogeneous progenitor cell pool and a single extracellular substance gradient that enables correct cell migration along the radial direction in 3D physical space. Hence, the phenotype is generated in a self-organizing way.

It is well-established that apoptosis is abundant during cortical development (Thomaidou et al. 1997; Cavallaro 2015). Already in the early days of neuroscience, the differential roles and mechanisms of apoptosis have been noted (Ernst 1926; Glücksmann 1951). Kuan et al. (2000) discuss the involvement of apoptosis, not only in cell number control, but also in the proper morphogenesis of the nervous system. Along those lines, Yeo and Gautier review regulatory mechanisms of neural programmed cell death (PCD) during vertebrate neural development (Yeo and Gautier 2004), which is different from neurotrophic death in differentiated neurons.

This study suggests that apoptosis during development comprises differential complexities; in an early apoptotic stage, cell death occurs independently from the local extracellular environment, and helps generate roughly the final cell numbers (A1-type). After layer formation has occurred, (A2-type) apoptosis serves a different role by improving the segregation between layers, hence reducing the overlap among layers. Here, the signals from neighboring cells that convey information on the suitability of cell positions within the extracellular context, are crucial. In our model, A2-type apoptosis is not necessary for the development of the layer architecture, but it improves the separation between layers. Notably, two distinct types of apoptosis have been observed experimentally (Rakic and Zecevic 2000), which could correspond to A1-type and A2-type apoptosis. It should be possible to experimentally verify the existence of these two hypothetical forms of apoptosis. For instance, A2-type apoptosis could be tested by displacing invidiual cells during critical developmental stages.

Importantly, the inclusion of A1-type apoptosis in our model enables a much wider variety of cortical layer architecture. If there was no such apoptosis, certain changes to the differentiation of early developing layers (e.g. to parameter C_6_ in Fig. 1) would necessarily impact all layers simultaneously, because these early differences influence the entire differentiation process later on. However, apoptosis during the entire developmental process grants more independence to the later developing layers to evolve differently from the earlier developing ones, because their apoptotic parameters also impact their final cell numbers. Hence, the number of cells in all layers can increase or decrease independently from other layers thicknesses, and apoptosis strongly improves this evolvability of the cortical layer architecture.

We highlight that certain observations naturally follow from our model, without tuning of model parameters. Along those lines, experimental work has demonstrated significant rates of apoptosis during proliferation (corresponding to A1-type apoptosis in our model) (Blaschke et al. 1996; Thomaidou et al. 1997). Indeed, also in our simulations the occurrence of A1-type apoptosis is usually comparable to the number of generated neurons (Suppl. Fig. S5). Moreover, the rate of embryonic cell death exhibited significant variation across time. This behaviour matches with the dynamics of our GRN model (Fig. 1), where apoptotic probabilities of the A1-type apoptosis depend on the GRN state. It is also in accordance with our model that apoptotic rates during development of homologous neuronal populations can vary significantly between species, as well as within the same species (Williams and Herrup 1988). Finally, it is established the crucial role that apoptosis plays in determining cortical thickness (Inglis-Broadgate et al. 2005).

Our model suggests that the timing of apoptosis is a crucial factor in the generation of the respective outcome, and so very different phenotypic properties in neurodevelopmental disorders can be generated depending on the time of apoptotic malfunction. This indicates that certain developmental brain disorders could in fact stem from causes that share many features but occur at different times. Indeed, a number of studies demonstrate the involvement of aberrant apoptosis in developmental brain disorders, such as in autism (Margolis et al. 1994; Wei et al. 2014), FASD (Creeley and Olney 2013), schizophrenia (Glantz et al. 2006) or microcephaly (Poulton et al. 2011). In addition to apoptosis, proliferative dynamics are crucial in generating the appropriate cell numbers and cell types (Lui et al. 2011). Nevertheless, distinct brain developmental disorders will affect proliferation and apoptosis to different extents, and can exhibit significant heterogeneity (Donovan and Basson 2017). Hence, the genetic causes can vary depending on the subtype of the developmental disorder, the specific brain region and the overall genetic context. Notably, specific changes to the proposed GRN model dynamics produce phenotypic outcomes that are in agreement with observations of healthy and pathological cortical tissue: changes to apoptosis early during development (A1-type) lead to thinner cortices (polymicrogyria), while introducing a defect to apoptosis later during development (i.e. A2-type apoptosis occurring after migration) gives rise to a more disorganized layer structure associated with autism. Migration deficits yielded pathological gray matter below deep cortical layers, in agreement with subcortical band heterotopia.

These findings are relevant from a clinical point of view. One common feature of developmental malformations is a difference in cortical thickness in various regions. Our model shows the importance of apoptosis in the control of appropriate layer formation and cortical cytoarchitecture. Given this central role of apoptosis, medical intervention to redirect apoptotic behaviour at suitable times during brain development is likely a fruitful direction for clinical research. Moreover, for substances that are administered during pregnancy or commonly used in early childhood, particular attention should be attributed to the apoptotic impact (Palanisamy 2012; Sinner et al. 2014). Indeed, reducing apoptosis leads to severe cortical malformation (Kuida et al. 1998). However, increased apoptosis is also a possible origin of disorder (Lainhart and Lange 2011).

Studies of polymicrogyria show increased apoptosis and a thinner cortex (Stottmann et al. 2013; Huang et al. 2016). We find that specific defects in gene-regulatory dynamics controlling apoptosis yield characteristic properties associated with this disorder. Intriguingly, the gene GPR56 has been associated with polymicrogyria (Bahi-Buisson et al. 2010) as well as apoptotic pathways (Ke et al. 2007; Yang and Xu 2012). Hence, our results are in agreement with the suggestion that polymicrogyria is not a disorder of neuronal migration (Judkins et al. 2011), and highlight the possibility that impaired apoptosis is a key player in the origins of certain neurodevelopmental disorders. However, in our simulations we introduced a single change to apoptosis occurring before cell migration is induced. Therefore, in contrast to Judkins et al. (2011), we do not suggest that polymicrogyria should be considered a post-migration malformation of cortical development. Instead, we propose that pathological apoptosis before and during migration could be the key driving cause.

Multiple studies demonstrate a tight link between local cytoarchitecture and global brain connectivity as defined by interregional, long-range fiber tracts (Hilgetag and Amunts 2016; Wei et al. 2018). In particular, regions of similar cytoarchitecture and laminar profile tend to be more likely connected. While there are multiple possible explanations for this relationship, it suggests that alterations of the cytoarchitecture due to neurodevelopmental disorders will also impact long-range connectivity changes. We anticipate that computational models bridging the local (Bauer et al. 2014) and global (Bauer and Kaiser 2017) spatial scales will help better understand such intricate developmental dynamics. Along those lines, we are developing software tools to overcome the challenges of reconciling such spatial heterogeneity (Bauer, Breitwieser, Di Meglio, Johard, Kaiser, Manca, Mazzara, et al. 2017; Gonzalez de Aledo et al. 2018; de Montigny et al. 2020).

Given that key neurodevelopmental genes are often involved in many functions (Manzini and Walsh 2011), a specific gene variant might operate correctly at certain developmental stages, but may be wrongly expressed at others. Future treatments will have to take into account the specific timing of genetic expression during development. Along those lines, we envisage that computational models of developmental processes will constitute an additional pillar for clinical research, allowing to anticipate fruitful experiments and formulate novel hypotheses.

Our proposed computational model gives rise to a number of experimentally verifiable and quantifiable predictions. Given the robustness of the core GRN structure across different species, apoptotic rates should be a major determinant of differences in layer-specific neuron numbers across species, as well as within the same species. This is in agreement with previous studies on human and hamster cortex, suggesting that apoptosis could have a key role in shaping regional differences (Finlay and Slattery 1983; Rakic and Zecevic 2000). Furthermore, our work suggests that it should be possible, by changing apoptotic events during brain development, to differentially impact cortical layers. In other words, the number of cells in individual layers should be controllable by transient changes to apoptotic rates during proliferation. Such relationship could be assessed experimentally by analysing markers of apoptosis at different developmental stages during cortical layer formation, in combination with layer-specific markers (Hevner 2007; Brunjes and Osterberg 2015). Finally, the computational model suggests that apoptosis, after cell migration has terminated, is relatively large for marginal zone cells. This is partially due the likelihood of not being pushed by later-developing layers, which is larger than for other layers (see Fig. 6). Additionally, in order to reproduce the small layer 1 neuron numbers, our model requires specifically high marginal zone cell apoptosis rates in the superficial layers. Indeed, experimental analyses are in agreement with frequent MZ cell apoptosis (Meyer and González-Gómez 2017).

From an evolutionary perspective, our work contributes to a better understanding of a longstanding conundrum on the question why there is so much apoptosis during brain development, i.e. approximately one-half of all neurons (Cavallaro 2015). One possibility is that it enables cell selection, i.e. inappropriate cells can be removed for the sake of a more refined cortical structure (Pompeiano et al. 2000). However, from an energetic point of view such massive apoptosis would be surprisingly inefficient. We propose that one key function of apoptosis during brain development is to render the neural architecture more evolvable, enabling the brain to adapt faster and more efficiently to different requirements. Along those lines, it has been previously suggested that apoptosis is responsible for the rapid evolutionary reduction in neuron populations in Spanish wildcats and their domestic relatives (Williams et al. 1993). Moreover, it is in accordance with this evolutionary perspective that apoptosis seems to operate in all multicellular animals (Ameisen 2002). Finally, apoptosis is known to enable greater flexibility as certain evolutionary old structures, such as for example pronephric kidney tubules, which form functioning kidneys in fish and in amphibian larvae, are not active in mammals and degenerate (Meier et al. 2000). As mentioned before, it is the appropriate balance between apoptosis and proliferation that is crucial to generate healthy neural structures. Our model suggests that human brain evolution was enabled by a suitable composition of proliferation that is in line with area-specific apoptosis, which determines the properties of the ontogenetic column (Geschwind and Rakic 2013).

We acknowledge that neuronal survival and apoptosis are also influenced by electrical activity (Heck et al. 2008) and axonal innervation (Popovik and Haynes 2000) but this third stage of apoptosis occurs outside of the timeframe of this simulation; cortical layers, other than marginal zone and subplate, are first established before the ingrowth of thalamic afferents, connection of corticofugal projections to their targets, and formation of intracortical networks (Huttenlocher and Dabholkar 1997; Kostovic 2002). As we do not model the formation of axonal branches and synaptic connectivity, it is beyond the scope of this work to cover the full developmental extent of apoptosis in cortex. Taking into account additional developmental mechanisms constitutes a key step towards an integrated study of physiological implications of normal and pathological brain development. Due to their important contribution to the cortical volume, modelling neurites and synapses will also improve accordance with measurements of neuronal density. First steps have been taken in this direction, leading to neuronal self-organization of biologically plausible axonal arborizations (Bauer et al. 2012), complex network properties of synaptic connectivity (Bauer and Kaiser 2017) and computationally powerful circuit function (Bauer et al. 2014). We also aim at extending our simulations using modern software tools (Gonzalez de Aledo et al. 2018; Bauer et al. 2017), which will enable simulations that give rise to a detailed model of neural tissue dynamics including folding patterns (Wang et al. 2016).

Given that our model can account for a wide range of layered neocortical cytoarchitecture, only small amendments are likely to be necessary to explain also other layered structures in the central nervous system, such as the retina or hippocampus. In the future, our model could be extended by integrating data on gene expression, such as obtained from public databases (e.g. (Sunkin et al. 2012; Lindsay et al. 2016)). Such detail will enable the modelling of the relationship between genetic specification and the associated phenotypic properties of cortex.

Overall, we provide a mechanistic, agent-based computational model for cortical layer formation that takes into account the interaction of intracellular dynamics with extracellular communication. In particular, our model implicates the role of apoptosis in enabling the evolvability and variety of cortical layer structure. Moreover, it constitutes a platform for the generation and testing of hypotheses, such as for example on the malfunctioning mechanisms that give rise to developmental disorders.

## Acknowledgement

The authors wish to thank Rodney Douglas, Rob Forsyth, Evelyne Sernagor, Peter Taylor and Yujiang Wang for stimulating discussions and useful comments. This work made use of the facilities of N8 HPC Centre of Excellence, provided and funded by the N8 consortium and EPSRC (Grant No.EP/K000225/1). The Centre is co-ordinated by the Universities of Leeds and Manchester. Moreover, parts of the computational work were performed using the “Rocket” HPC at Newcastle University.

