## Supplemental Information for "Creative destruction: a basic computational model of cortical layer formation"

### Supplementary information for "Dying to evolve: a basic computational model of cortical layer formation"

#### 1 Layer formation

The video Suppl. Video S1 shows a simulation of layer formation in human temporal cortex (Fig. 3). An initial, homogeneous precursor pool of proliferating cells gives rise to different neuron types in a given order. Each cell comprises an individual gene-regulatory network, and interacts with its local external environment. The final cortical column comprises approximately 7,000 neurons. The video can also be found on Youtube: `https://youtu.be/pYxnK3YXQmU`.

#### 2 Normalised layer-specific neuron distribution

Suppl. Fig. S1 shows the simulated distribution of neurons across different neocortical layers.

#### 3 Dependence of layer architecture for examplary parameter variation

Suppl. Fig. S2 shows the impact of changing examplary GRN parameters on the layer architecture.

#### 4 Polymicrogyria

We conducted simulations of a pathological neurodevelopmental scenario, where cortex is thinner than in the control case. As shown in the manuscript, such a thinning can be obtained by increasing apoptosis during the first apoptotic stage (states A1 in Fig. 1, see also Suppl. Fig. S2). Alternatively, the exponential proliferation phase can be shorted by increasing the threshold for initiation neural differentiation. In this case, the early proliferative phase consisting of symmetric division is shortened, hence fewer neurons are generated.

#### 5 Subcortical heterotopia

We also simulated the developmental disorder of subcortical band heterotopia, where heterotopic neurons in the proliferative zone can be observed (Suppl. Fig. S4). Introducing defects to neuronal migration gives rise to characteristics of malformation in agreement with this disorder.

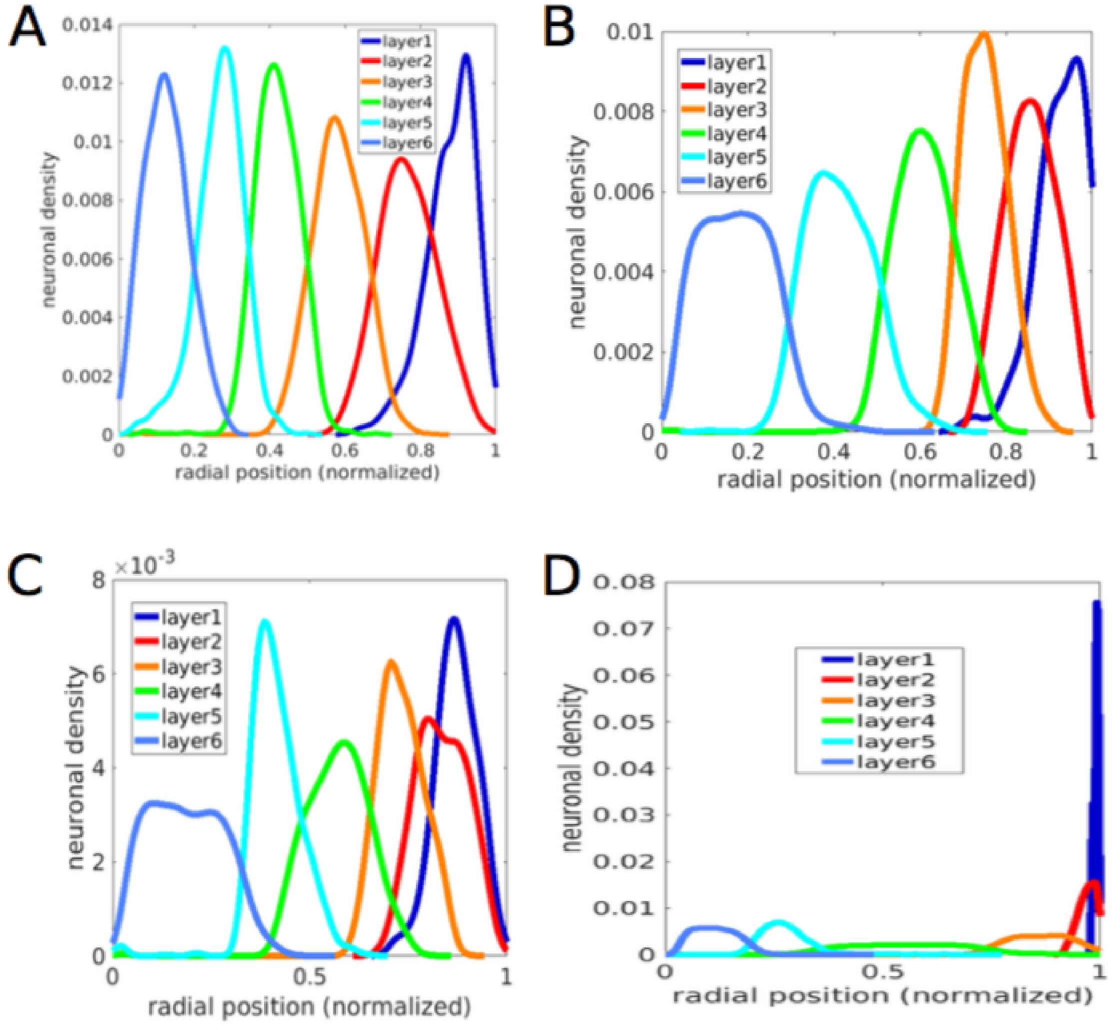


Figure S1: Normalised densities of neurons in different layers. This figure complements Fig. 4, which visualizes the distributions of the number of neurons across the radial direction.

#### 6 Differential modes of apoptosis

Our model incorporates two modes of apoptosis, i.e. type A1 and type A2. Type A1 is independent on environmentanl signals, while type A2 depends on the cell types in the local neighborhood of cells. Suppl. Fig. S5 shows examplary numbers of occurrences of apoptosis in a simulation of layer formation in human temporal cortex. In agreement with experimental data, apoptosis is highly abundant during cortical development.

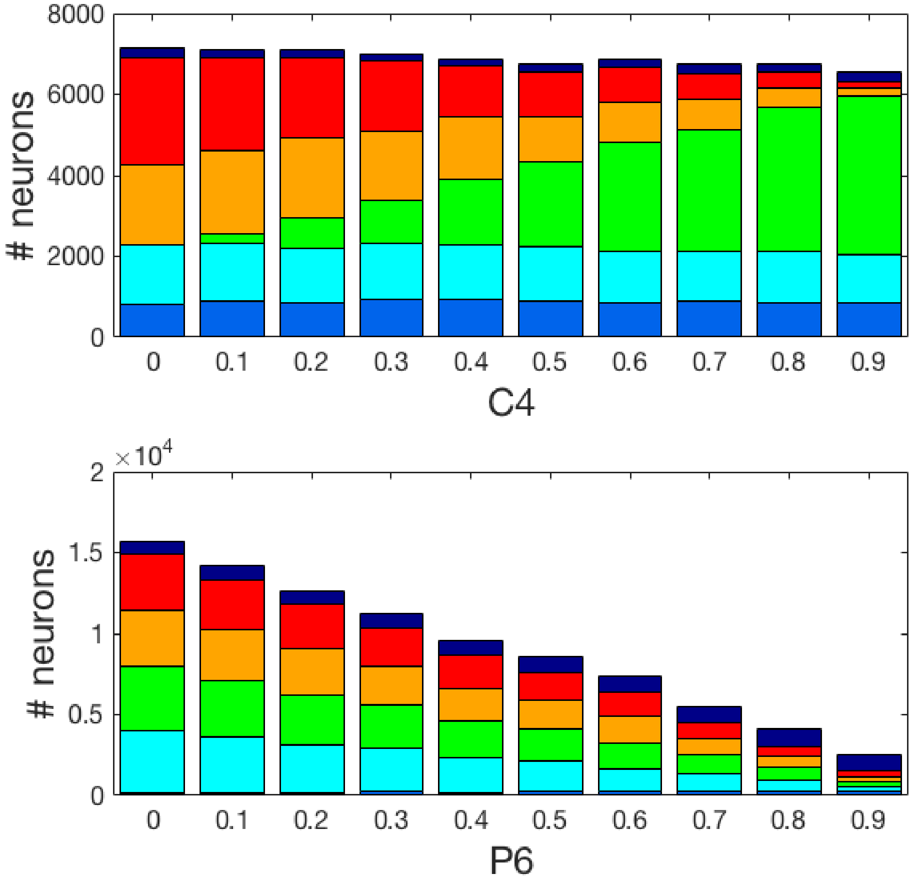


Figure S2: Layer architectures for different gene regulatory network parameter settings. In the upper plot, the probability to commit to layer 4 is varied, showing its impact on the number of neurons of layers 4, 3 and 2. In the bottom plot, the apoptotic probability P6 is varied, influencing the number of neurons across all layers, due to the early activation of this apoptotic process.

### 7 Experimental Data

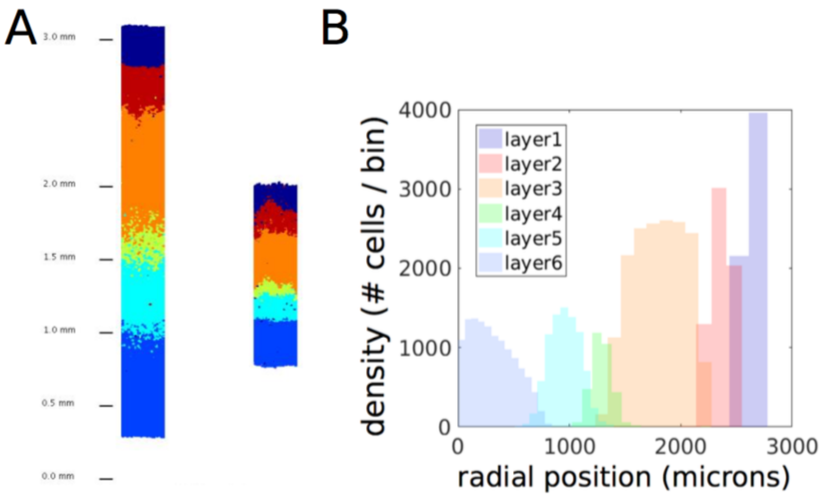


Figure S3: Impact of shorter proliferation phase on cortical layer architectures. (A) Left: Control layer architecture, in agreement with layer thickness measurements from human peristriate area 19. Right: Reduced cortical thickness due to the exponential proliferation phase terminating prematurely (while the differentiation phase is not altered but induced earlier). (B) Histogram of neuronal locations in the control layer architecture from (A).

Table S1: Estimated layer-specific numbers of neurons across different species, using experimental data from O'Kusky and Colonnier 1982 (for human, rat and mouse data) and DeFelipe et al. 2002 (for macaque data).

| #neurons under $1mm^2$ in | Human temporal cortex | Rat hindlimb somatosensory cortex | Mouse barrel cortex | Macaque visual cortex |
|---|---|---|---|---|
| Layer 1 | $1,958$ | 427 | $1,257$ | 600 |
| Layer 2/3 | $27,514$ | $28,183$ | $32,346$ | $46,199$ |
| Layer 4 | $13,158$ | $13,827$ | $37,723$ | $90,300$ |
| Layer 5 | $12,738$ | $20,785$ | $19,286$ | $24,800$ |
| Layer 6 | $8,051$ | $36,322$ | $55,063$ | $30,199$ |

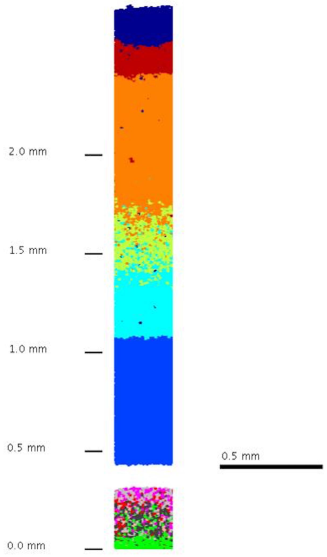


Figure S4: Simulation of cortical organization in subcortical heterotopia. Characteristic features of subcortical heterotopia or double-cortex syndrome are recapitulated by introducing migration defects. Here, 30 % of differentiated neurons exhibited pathological migration and remained in the proliferative zone. As a consequence, heterotopic neurons are observed below the deep cortical layers, while cortical layer structure is preserved.

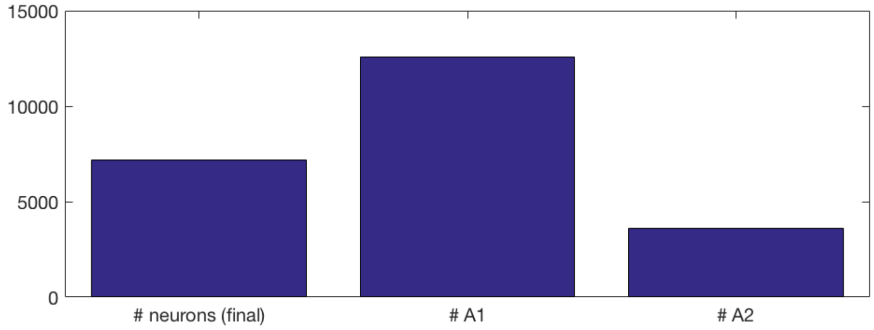


Figure S5: This figure shows the final number of neurons after simulation of layer formation in human temporal cortex (left), the number of cells that have died during development due to A1-type apoptosis (middle), and the number of neurons that have died due to A2-type apoptosis. In accordance with experimental observations, the occurrence of apoptosis during cortical development is in the same order of magnitude as the number of surviving neurons.
